# Insights into the molecular mechanisms underlying the inhibition of acid-sensing ion channel 3 gating by Stomatin

**DOI:** 10.1101/704064

**Authors:** Robert C. Klipp, Megan M. Cullinan, John R. Bankston

## Abstract

Stomatin is a monotopic integral membrane protein found in all classes of life that has been shown to regulate members of the Acid-Sensing Ion Channel (ASIC) family. However, the mechanism by which Stomatin alters ASIC function is not known. Using chimeric channels, we combined patch clamp electrophysiology and FRET to search for regions of ASIC3 critical for binding to and regulation by Stomatin. With this approach, we found that regulation requires two distinct sites on ASIC3: the distal C-terminus and the first transmembrane domain. Mutation of the C-terminal site disrupts binding and regulation whereas disruption of the transmembrane site eliminates functional regulation. We then showed that Stomatin does not alter surface expression using fluorescence imaging and a surface biotinylation assay. Based on these findings, we propose a model whereby STOM is anchored to the channel via a site on the distal C-terminus but alters ASIC3 gating through action on TM1.

## Introduction

The Acid-Sensing Ion Channels (ASICs) are members of the Degenerin/Epithelial Na^+^ channel (DEG/ENAC) family of ion channels. ASICS are Na^+^ selective, voltage-insensitive channels that are activated by extracellular protons. Six mammalian ASIC isoforms are known to exist, which can form either heteromeric or homomeric trimers (Hesselager et al., 2004; Jasti et al., 2007). ASIC1a, 1b, 2a, and 3 all form functional pH sensitive channels as homotrimers, while ASIC2b and 4 homotrimers are not gated by protons (Akopian et al., 2000; Gründer et al., 2000; Lingueglia et al., 1997). ASICs are expressed throughout the central and peripheral nervous systems where they are thought to play a role in a range of physiological and pathophysiological functions including nociception, fear conditioning, neuronal death following ischemia, baroreception and autonomic control of circulation, and sensing myocardial ischemia (Benson et al., 1999; Jones et al., 2004; Lu et al., 2009; Ugawa et al., 2002; Wemmie et al., 2002; Xiong et al., 2004). Like many of the ion channels of the sensory system, ASICs are multimodal receptors; in addition to activation by protons, ASICs are regulated by lipids, phosphorylation, numerous extracellular ligands, and accessory proteins (See review (Boscardin et al., 2016)).

A number of high-resolution structures of ASIC1a from chicken have been solved (Baconguis et al., 2014; Baconguis and Gouaux, 2012; Dawson et al., 2012; Jasti et al., 2007; Yoder et al., 2018). These structures have provided hypotheses for how protons and toxins derived from animal venoms might act on the extracellular domain of the channel and lead to opening and closing of the gate. However, in each structure, the intracellular domains are either missing from the protein or not resolved in the structure. Consequently, little is known about how the intracellular termini contribute to channel function and how proteins that interact in this region might impact channel function.

Stomatin (STOM) is a 31.5 kDa monotopic integral membrane protein ubiquitously found throughout the central and peripheral nervous system. STOM is associated with the cytoplasmic face of the plasma membrane via a hydrophobic hairpin region and a number of palmitoylation sites (Snyers et al., 1999). In humans, there are four closely related proteins to STOM which are the Stomatin-like proteins 1-3 (STOML1, STOML2, STOML3) and a protein important for proper filtration in the kidney called Podocin (See review (Browman et al., 2007)). In addition, STOM is part of a superfamily of proteins that contain a conserved Stomatin, Prohibitin, Flotillin, HflK/C domain (SPFH) domain. STOM has previously been shown to regulate several membrane proteins including: the Glucose Transporter GLUT-1, the Anion Exchanger AE-1, and ASICs (Brand et al., 2012; Genetet et al., 2017; Moshourab et al., 2013; Price et al., 2004; Zhang et al., 2001).

Previous work showed that recombinant expression of STOM into mammalian cells decreased ASIC3 current magnitude, sped ASIC2a desensitization, but had no effect on ASIC1a (Price et al., 2004). Additionally, other members of this family of proteins including the STOMLs, Podocin, and the Flotillins have all been shown to regulate other ion channels as well (Anderson et al., 2013; Kozlenkov et al., 2014; Lapatsina et al., 2012; Liu et al., 2016; Moshourab et al., 2013; Wetzel et al., 2007). STOMLs in particular have been shown to regulate ASICs in an isoform dependent manner. The STOM homologue, MEC-2 is essential for the function of the mechanosensitive ASIC homolog MEC-4/MEC-10 complex in *Caenorhabditis elegans* (Brown et al., 2008; Goodman et al., 2002; Huang et al., 1995). Despite the ubiquity of this family of proteins, little is known about the mechanisms through which STOM and its family members regulate ion channels.

Here, we paired patch clamp electrophysiology with Förster Resonance Energy Transfer (FRET) to localize the binding site for STOM on ASIC3. Making chimeric channels between ASIC3, which is regulated by STOM, and ASIC1a, which is not regulated, allowed us to localize two sites on ASIC3 that are critical for STOM-dependent regulation. First, we found that the distal C-terminus of the channel was necessary for both complex formation and regulation of the channel. Second, the first transmembrane domain (TM1) of ASIC3, while not sufficient for binding, was necessary for STOM-dependent regulation. In addition, we showed that STOM does not act by altering the cell surface expression of ASIC3 and likely alters function by changing channel gating. Taken together this led us to a model whereby STOM binds to the distal C-Terminus of ASIC3 but largely exerts its effect through a second interaction with TM1 of the channel. These results extend our understanding of the STOM/ASIC3 complex and may shed light on how the SPFH domain proteins regulate ASICs more generally.

## Results

### Stomatin binds to ASIC3 and dramatically inhibits total current without reducing channel surface expression

Work on STOM and the STOML family of proteins has suggested that these proteins regulate ASIC gating in an isoform dependent manner (Kozlenkov et al., 2014; Lapatsina et al., 2012; Moshourab et al., 2013; Price et al., 2004). STOM is a 31.5 kDa integral monotopic protein with several interesting structural features (Fig. 1A). 1) STOM has a short N-terminus and a small hairpin that is important for membrane localization. 2) STOM contains an SPFH domain. 3) STOM has multiple palmitoylation sites, at least one of which is required for membrane targeting (Snyers et al., 1999). 4) STOM contains at least one cholesterol binding site and is frequently localized to cholesterol-rich lipid rafts. Given the topology of STOM, we reasoned that the binding site for STOM on ASIC3 must involve the transmembrane domains and/or the intracellular termini.

**Figure 1.**
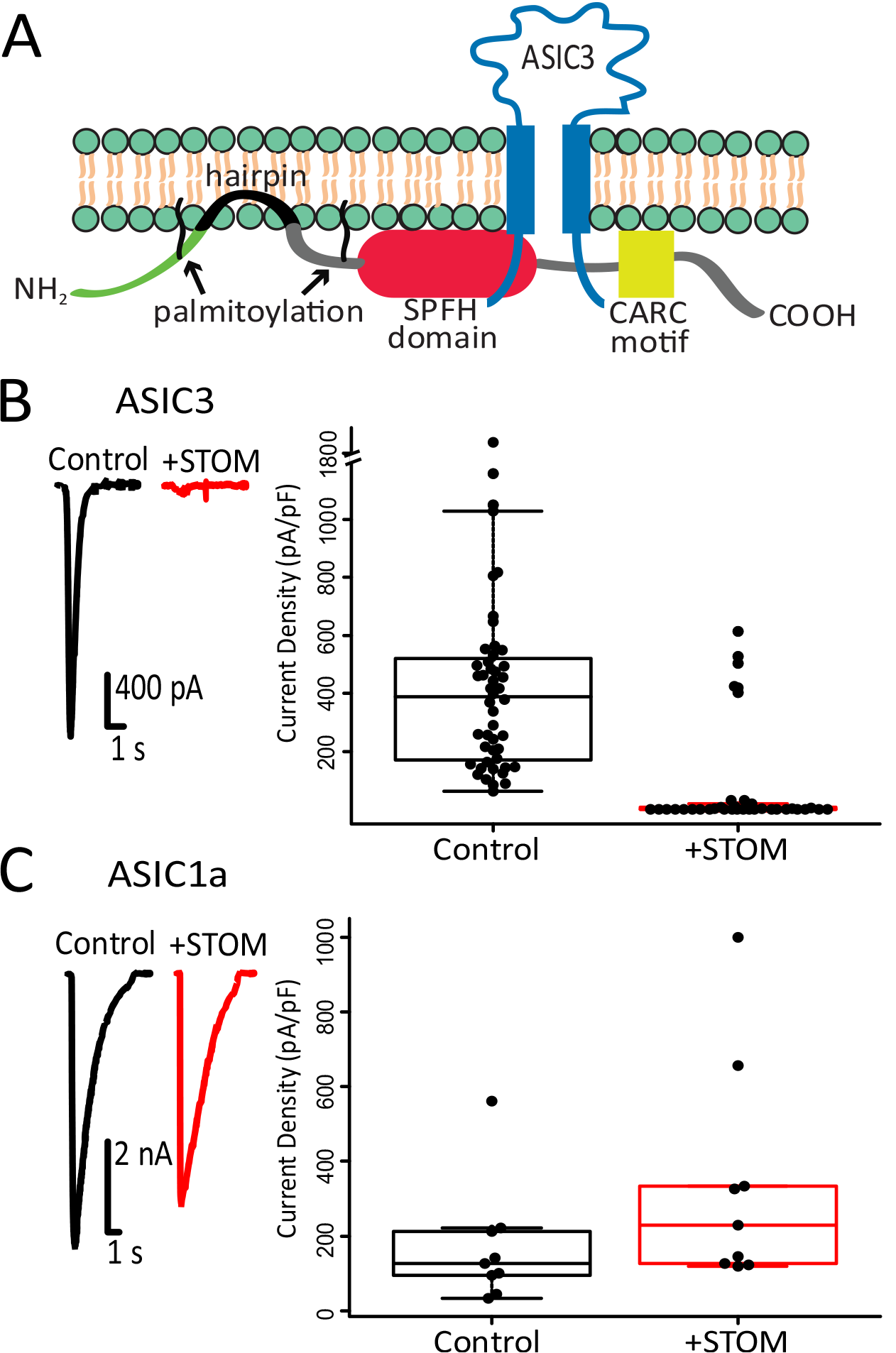
STOM inhibits ASIC3 but not ASIC1a current amplitude. (**A**) Topological cartoon of STOM interaction with ASIC3. (**B**) Left shows representative pH 5.5 evoked ASIC3 currents alone (black) and co-transfected with STOM (red). Plot to the right shows every measurement made for the 2 conditions plotted as the current density (Peak current amplitude/Cell capacitance) superimposed onto a boxplot that summarizes the data. Average current density of the control and +STOM conditions were 364.0 ± 32.6 pA/pF (n=45) and 2.0 ± 0.4 pA/pF (n=27), respectively. (**C**) Identical experiments as in panel B performed for ASIC1a demonstrates that co-expression with STOM (red) does not inhibit acid-evoked currents. Average current densities of ASIC1a in the absence (black) and presence (red) of STOM were 122.5 ± 22.9 pA/pF (n=8), and 200.3 ± 33.7 pA/pF (n=7), respectively. All data is given as Mean ± SE.

Previous reports have suggested that STOM dramatically reduces ASIC3 currents and that this change in ASIC3 current does not occur due to a change in cell-surface expression (Price et al., 2004). We first sought to confirm these original findings. We used patch clamp electrophysiology to measure the functional effect of STOM co-expression with ASIC3 in CHO-K1 cells. Whole-cell patch clamp recordings were performed by rapidly switching between solutions at pH 8.0 and pH 5.5 using a piezo-driven solution exchange system. Representative current traces in Figure 1B, shows that control ASIC3 acid evoked currents (black) are drastically reduced in the presence of STOM (red). Boxplot summary of all measurements made, also plotted in Figure 1B, shows that STOM reduced ASIC3 mean current density from 364.0 ± 32.6 pA/pF (n=45) to 1.96 ± 0.44 pA/pF (n=27)). Performing the same experiment for cells expressing ASIC1a with and without STOM also confirmed previous findings that ASIC1a was not functionally regulated by STOM (Fig. 1C). In 6 out of the 35 recordings, ASIC3 displayed control-like current densities even in the presence of STOM. This is likely an artifact of our transient transfection system where STOM may be either absent or weakly expressing in these cells. Given these outliers, we calculated the mean current density for every experiment in this study in two ways. First, we averaged all the data collected. Second, we excluded outliers that were in excess of 1.5*IQR, where IQR is the inter-quartile range of the data. We will use this adjusted mean to discuss the data, but both calculations appear in Supplemental Table 1. Additionally, all data, including outliers, are shown throughout in the boxplots.

We next sought to determine if a change in surface expression could explain the dramatic reduction in ASIC3 currents when STOM was present. A change in channel current density must occur either via a change in channel gating, or a change in surface expression. We elected to examine membrane expression of ASIC3 in two ways. First, we used confocal microscopy to look at the localization of fluorescently tagged ASIC3 to determine if STOM co-expression dramatically altered the distribution of channels in the cell. To do this, we co-expressed ASIC3 with a C-terminal Turquoise tag (ASIC3-TUR) and STOM with a C-terminal YFP tag (STOM-YFP). To help ensure that we could identify the plasma membrane, we used a TagRFP labelled portion of the L-type calcium channel, consisting of the I-II loop of Ca_v_1.2, joined to the N-terminus of Ca_v_1.1. This protein, which we will designate “membrane marker” in the figure, has previously been shown to associate with the plasma membrane (Kaur et al., 2015; Polster et al., 2018). Figure 2A shows representative images with and without co-expressed STOM-YFP. In each case, ASIC3-TUR was strongly associated with the cell surface. Figure 2B shows line-scans of both the RFP and TUR signals both with and without STOM-YFP. Peaks in fluorescence intensity for ASIC3-TUR coincided with peaks from our RFP membrane marker suggesting that ASIC3-TUR was on the plasma membrane. This membrane localization was not altered by STOM co-expression.

**Figure 2.**
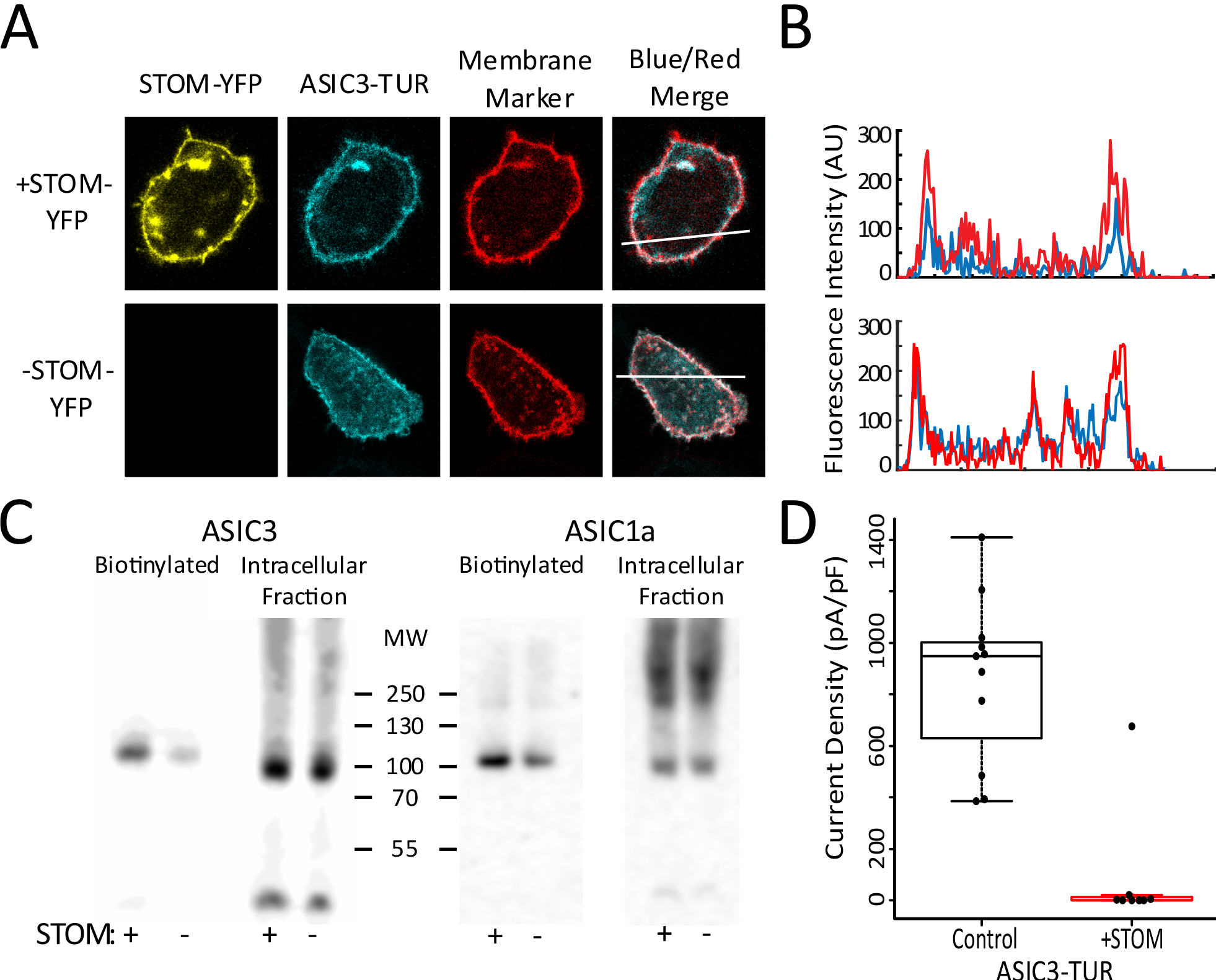
STOM does not alter ASIC3 expression on the plasma membrane. (**A**) Confocal images of cells expressed with ASIC3-TUR, membrane marker (TagRFP labelled I-II loop of Ca_v_1.2 with Ca_v_1.1 N-terminus), with (Top) and without STOM-YFP (Bottom). (**B**) Line scan of the Blue and Red signals from (A) (indicated by white bar) shows that ASIC3-TUR peak intensity overlaps with membrane marker on the plasma membrane in the presence and absence of STOM-YFP. (**C**) Cell surface biotinylation of ASIC3-TUR and ASIC1a-CER with and without STOM indicates that STOM does not decrease biotinylated (plasma membrane) ASIC3-TUR or ASIC1a-CER surface expression. (**D**) Boxplot showing the current density measured from pH 5.5 evoked currents from ASIC3-TUR with and without STOM confirm that fluorescently-tagged ASIC3 construct used in biotinylation assay displays normal STOM inhibition. Current density for control and +STOM were 859.2 ± 94.1 pA/pF (n=11) and 4.6 ± 2.7 pA/pF (n=7), respectively. Data given as Mean ± SE.

As a second method for examining membrane expression, we employed a cell-surface biotinylation assay where we expressed the same ASIC3-TUR with and without untagged STOM. One day after transfection of our proteins into CHO-K1 cells, we isolated plasma membrane localized proteins by labelling them with Sulfo-NHS-SS-Biotin and pulling them down with NeutrAvidin resin. We then performed Western blots looking at the surface membrane and intracellular fractions of ASIC3. To ensure equal loading of protein in our Western blots, we quantified total protein concentration in our lysate using a BCA assay and adjusted our gel loading to ensure that we loaded equal amounts of total protein into each lane. The blot in Figure 2C shows that co-expression of STOM did not dramatically reduce the total amount of ASIC3 on the membrane. In fact, there appeared to be a modest increase in total ASIC3 protein on the membrane. In this example, there was a 97% increase in ASIC3 on the plasma membrane when STOM was co-expressed. On average, we found a 37 ± 31.8% increase in ASIC3 surface expression when STOM was present (n=3) with only a 3% change in total ASIC3 expression. To ensure these results were not impacted by the presence of the fluorescent protein on the C-terminus of ASIC3 we measured whole-cell acid evoked currents of ASIC3-TUR with and without STOM. We found that STOM was able to dramatically inhibit ASIC3-TUR to the same extent as the untagged channel (Fig. 2D). ASIC1a surface expression also appeared to be modestly increased by co-expression with STOM. These data are consistent with our functional measurements that showed a trend towards larger currents when STOM was present in the case of ASIC1a (Fig. 1C). This modest increase in surface expression of both ASIC1a and ASIC3 when STOM was co-expressed may or may not be significant, but the data show clearly that the nearly 200-fold decrease in ASIC3 current when STOM was present cannot be explained by a change in cell surface expression.

### FRET shows a direct interaction between STOM and ASIC3

We next wanted to develop an approach that would allow us to measure direct interaction between ASIC3 and STOM. To do this, we used acceptor photobleaching Förster Resonance Energy Transfer (FRET). We tagged ASIC3 with a C-terminal Cerulean (CER) and STOM with a C-terminal YFP and recorded two cyan (in response to weak 458 nm excitation) and two yellow (in response to weak 514 nm excitation) images. Subsequently, the cell was subjected to repeated illuminations with high intensity 514 nm light, which bleached YFP but had no effect on the CER. Finally, two more images were measured in each color using the same conditions as prior to bleaching the YFP. Because of the near total bleaching of YFP, if the two fluorophores are within ~70 nm of each other, the post-bleaching CER signal will be larger due to loss of resonant energy being donated to the YFP. A set of images, one before and one after bleaching, illustrating this method can be seen in Figure 3A. Using this approach, we measured an average FRET efficiency between ASIC3-CER and STOM-YFP of 12.5 ± 1.2% (n=11) (Fig. 3B) suggesting that these two proteins were interacting. Previous reports had suggested that STOM could bind ASIC1a as well, but our FRET assay did not detect interaction between the two molecules (Fig. 3B). Cells co-expressing only free CER and YFP showed no change in CER intensity between the pre- and post-bleaching images (data not shown). To ensure bleaching of the CER signal was minimal during the measurement we used very low laser power to excite the CER. Overall, the average decrease in CER intensity from the first to the final image in cells where only ASIC3-CER was expressed was only 0.8 ± 0.3%. This suggested to us that bleaching made only a minimal contribution to the results. With these data, we could now pair our functional measurement with our FRET measurement to attempt to localize sites on ASIC3 that are critical for binding STOM and sites that are critical for the functional regulation of ASIC3 by STOM.

**Figure 3.**
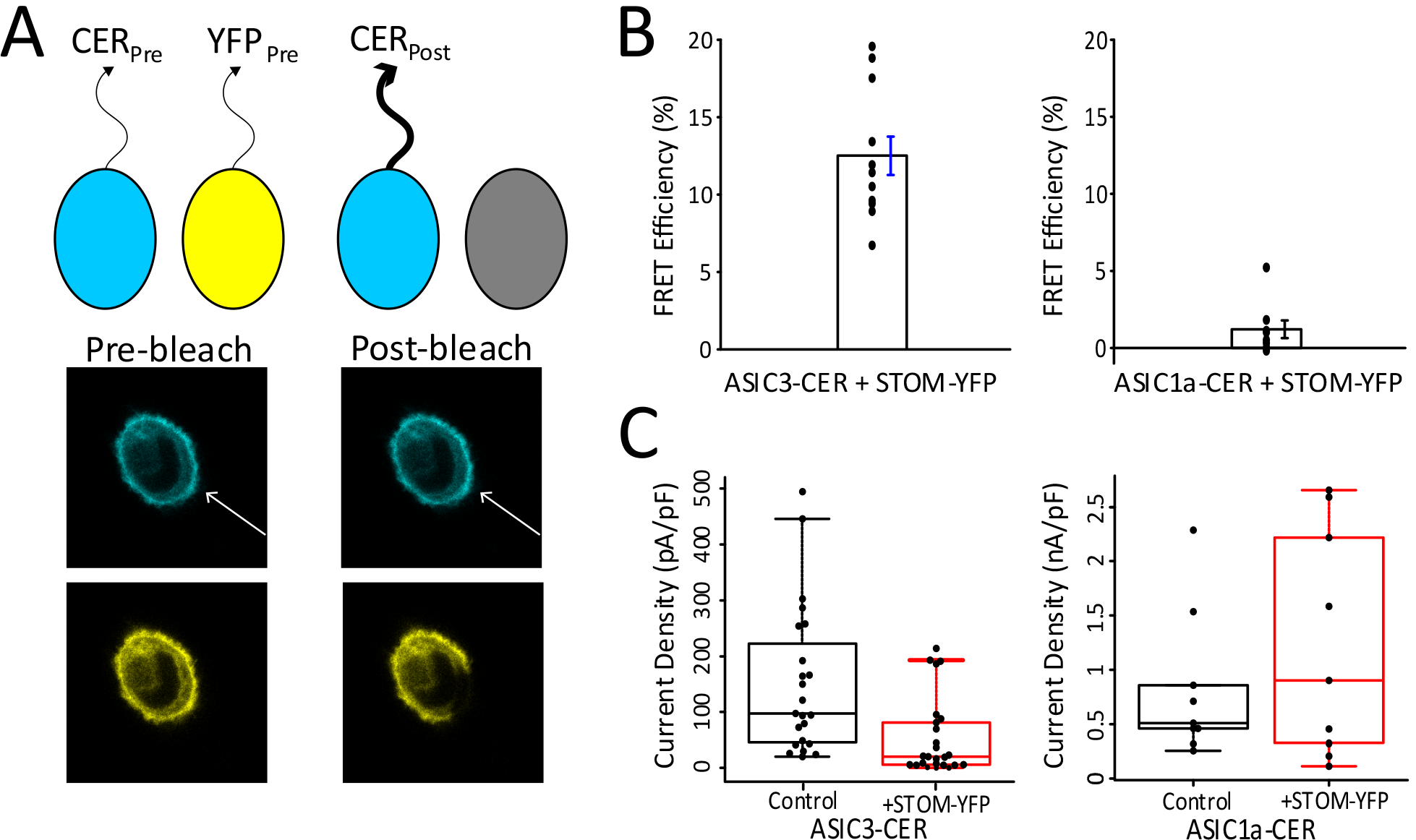
STOM inhibition of ASIC3 corresponds to a direct interaction. (**A**) Cartoon showing FRET photobleaching assay. Bottom panel shows a representative cell where the YFP was bleached which led to a corresponding increase in CER intensity. (**B**) Summary of FRET data showed that an interaction was measured between ASIC3-CER and STOM-YFP (Left) but not ASIC1a-CER (Right). FRET efficiency for ASIC3-CER and ASIC1a-CER were 12.5 ± 1.2% (n=11) and 1.2 ± 0.6% (n=8), respectively. (**C**) Boxplots showing the current density measurements for ASIC3-CER and ASIC1a-CER with and without STOM-YFP. Average current densities of ASIC3-CER in the absence (black) and presence (red) of STOM-YFP were 137.0 ± 23.5 pA/pF (n=22) and 47.1 ± 12.4 pA/pF (n=24), respectively. Average current densities of ASIC1a in the absence (black) and presence (red) of STOM-YFP were 509.7 ± 74.2 pA/pF (n=7) and 1227.5 ± 329.8 pA/pF (n=9), respectively. Data given as Mean ± SE.

To confirm that our fluorescent labels did not prevent STOM inhibition, we also carried out functional assays identical to those in Figure 1 using our fluorescently labelled proteins. Although inhibition of ASIC3 was maintained, we did observe that the magnitude of STOM inhibition of ASIC3 was diminished in the presence of the fluorophores (Fig. 3C). Compared to the control, mean current density decreased approximately 3-fold when ASIC3-CER was expressed with STOM-YFP, and again ASIC1a currents were not affected by STOM (Fig. 3C). We believe this reduced STOM effect occurs due to the YFP tag on STOM because the tagged ASIC3 in figure 2D was fully regulated by an untagged STOM and because STOM-YFP also showed a reduced inhibition of untagged ASIC3 (Fig. 3 - Supplement 1A). The YFP on the C-terminus of STOM could reduce the effect on ASIC3 in several ways. First, the STOM-YFP may show lower expression in our transient transfection system. Second, the presence of the fluorophore on the C-terminus of the channel allowed for us to select the brightest cells in each of the control and +STOM-YFP cases. This may have caused us to select cells where there was not enough STOM-YFP to fully regulate all of the ASIC3 in the cells. Consistent with each of these ideas, it appears from our data that cells with higher levels of STOM-YFP can inhibit ASIC3-CER as well as in the untagged case (Fig. 3 - Supplement 1B). Lastly, the YFP may simply interfere with the ability of STOM to regulate ASIC3. Despite this, STOM-YFP clearly interacted with ASIC3-CER and significantly reduced currents.

### Stomatin inhibition of ASIC3 requires the C-terminus

Having measured a direct interaction between ASIC3 and STOM, we then sought to identify the specific sites on ASIC3 that are critical for STOM regulation. To do this, we systematically created chimeras where portions of ASIC1a replaced the corresponding residues of ASIC3. With these chimeric channels we measured both the functional effect of co-expressing STOM as well as FRET between STOM and ASIC3. This allowed us to look for sites on ASIC3 that when mutated altered binding and regulation and sites that only altered the functional effect of STOM. For these first experiments, we used the untagged versions of both the channel and STOM because of the large effect and then used the tagged versions for the FRET measurements. We focused on domains in ASIC3 capable of directly interacting with STOM. Since STOM is a cytoplasmic monotopic integral membrane protein associated with the inner leaflet of the plasma membrane, potential interaction sites included the transmembrane domains and intracellular termini of ASIC3. Our chimera naming scheme follows the example in figure 4A, where we first indicate the isoform backbone followed by the backbone’s residue numbers being swapped for the corresponding residues of the other ASIC isoform. The protein alignment and domains are given in figure 4 - Supplement 1.

**Figure 4.**
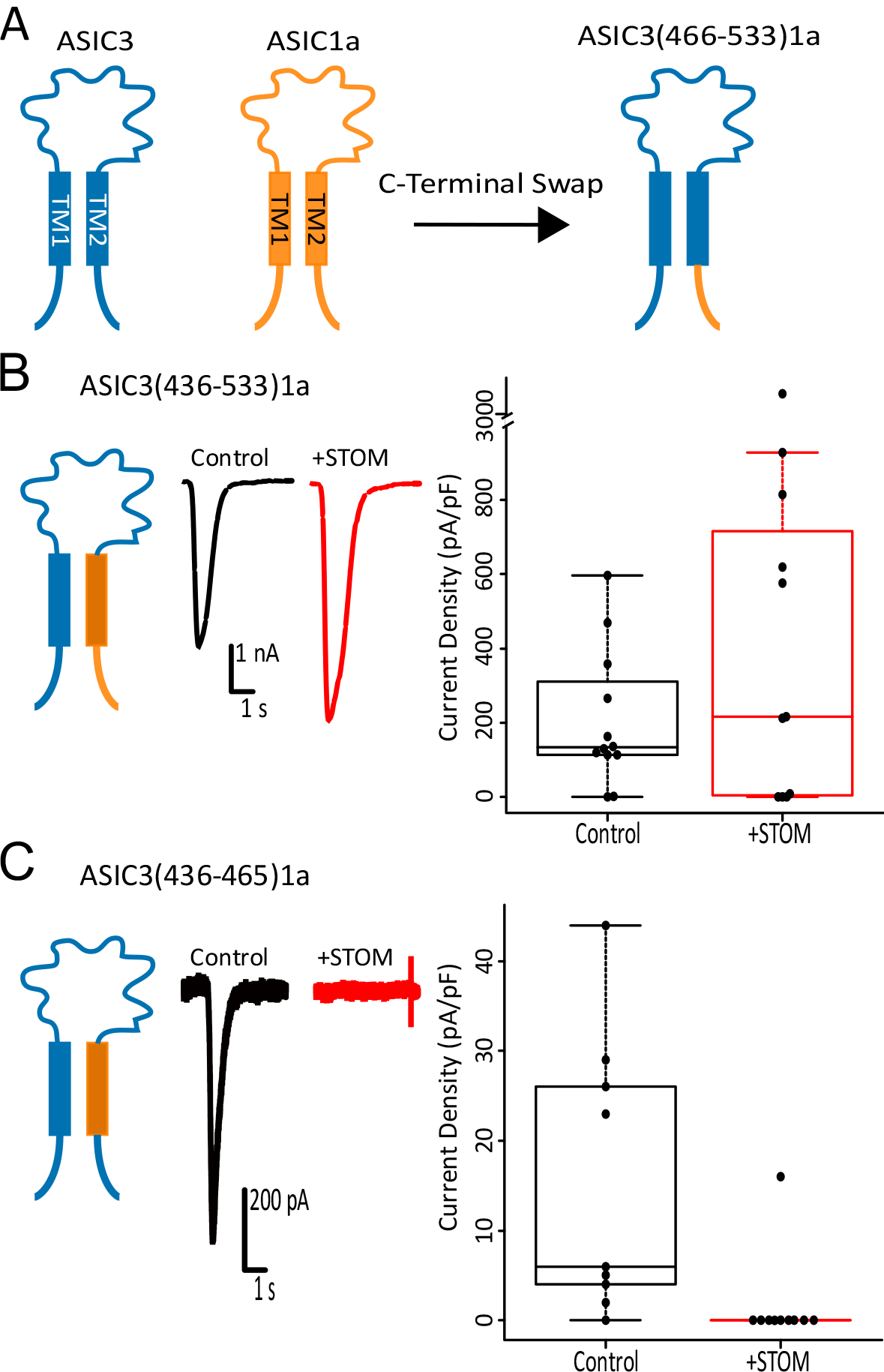
STOM inhibition of ASIC3 does not involve TM2. (**A**) Cartoon showing swap of ASIC3’s C-terminus (residues 466-533) with ASIC1a’s C-terminus (residues 459-526) is named ASIC3(466-533)1a. (**B-C**) Left shows cartoon depicting chimeras being examined. Middle shows representative pH 5.5 evoked currents of the indicated chimeric channel alone (black) and co-transfected with STOM (red). Right shows boxplots of the current density measurements for the chimeric channel with and without STOM. Swap of the TM2 and C-terminus of ASIC3 eliminates functional inhibition by STOM (B), but replacement of only TM2 of ASIC3 with the corresponding region of ASIC1a did not alter STOM-dependent regulation of ASIC3 (C). Average current densities for ASIC3(436-533)1a in the absence (black) and presence (red) of STOM were 205.3 ± 50.6 pA/pF (n=12) and 337.1 ± 109.0 pA/pF (n=10), respectively. Average current densities for ASIC3(436-465)1a in the absence (black) and presence (red) of STOM were 15.4 ± 4.9 pA/pF (n=9), STOM mean 0 ± 0 pA/pF (n=9), respectively. Data given as Mean ± SE.

We first investigated the C-terminus and transmembrane 2 (TM2) domains of ASIC3. Figure 4B shows that a swap of TM2 and the C-terminus of ASIC3 with the corresponding residues of ASIC1a, ASIC3(436-533)1a, eliminated STOM’s functional inhibition suggesting that this region is necessary for regulation. Replacement of TM2 alone, ASIC3(436-465)1a, resulted in a chimeric channel that expressed poorly. However, when we co-expressed STOM there was a clear and dramatic reduction in current suggesting that TM2 likely does not play a role in STOM-dependent regulation of ASIC3 (Fig. 4C).

Together these data suggested that the C-terminus is a critical determinant of the STOM-dependent regulation of ASIC3. To test this, we broke down the C-terminus of ASIC3 further. Replacing the entire ASIC3 C-terminus with that of ASIC1a, ASIC3(466-533)1a, eliminated STOM’s functional inhibition of ASIC3 (Fig. 5A). FRET measurements between ASIC3(466-533)1a-CER and STOM-YFP indicated that this loss of functional interaction was associated with loss of binding (Fig. 5A). Further breakdown of this region indicated that replacement of the first 40 residues of the C-terminus of ASIC3, ASIC3(466-505)1a, did not disrupt STOM inhibition or interaction with ASIC3 as measured by FRET (Fig. 5B). However, swap of the final 28 residues of the C-terminus, ASIC3(506-533)1a eliminated STOM inhibition as well as STOM binding to ASIC3 (Fig. 5C). These data point to the distal C-terminus as a critical determinant for the function and interaction of the STOM/ASIC3 complex.

**Figure 5.**
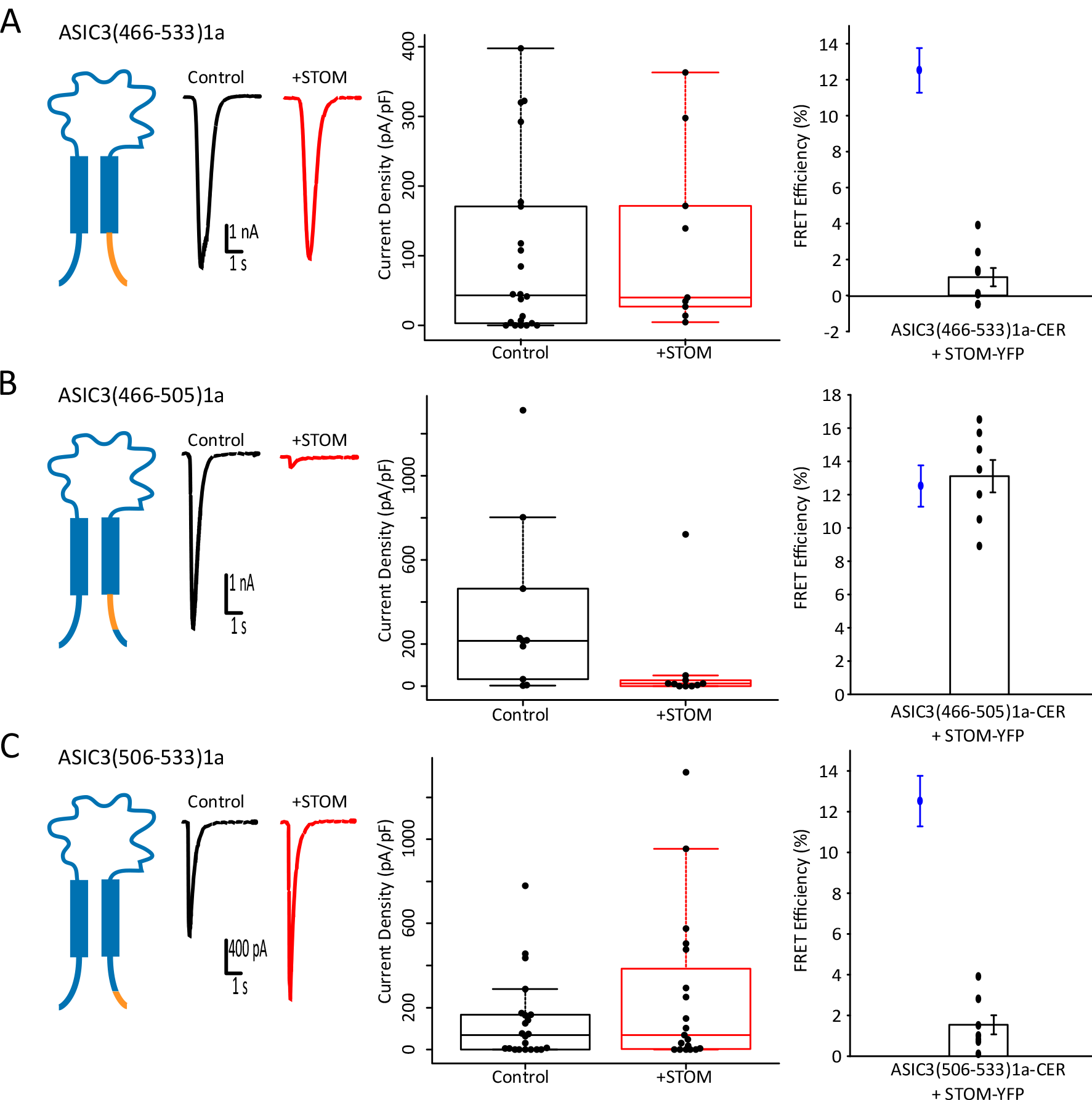
STOM binding at the distal C-terminus of ASIC3 is necessary for inhibition. (**A-C**) Left shows cartoon depicting the chimera being examined and representative pH 5.5 evoked currents of the indicated chimeric channel alone (black) and co-transfected with STOM (red). Middle shows boxplots of the current density measurements for the chimeric channel with and without STOM. Right panel shows the FRET efficiency between STOM-YFP and the indicated ASIC3-CER chimera. Replacement of the full C-terminus of ASIC3 (A) or the final 28 amino acids of the C-terminus (C) with the corresponding residues of ASIC1a eliminated STOM-dependent regulation of ASIC3 as well as binding. However, swap of only the first 40 amino acids (B) did not disrupt the interaction between ASIC3 and STOM nor the functional regulation. Average current densities for ASIC3(466-533)1a in the absence (black) and presence (red) of STOM were 99.5 ± 26.2 pA/pF (n=22) and 121.4 ± 41.7 pA/pF (n=9), respectively. Average FRET efficiency = 1.0 ± 0.5% (n=8). Average current densities for ASIC3(466-505)1a in the absence (black) and presence (red) of STOM were 239.9 ± 80.3 pA/pF (n=9) and 13.4 ± 5.1 pA/pF (n=9), respectively. Average FRET efficiency = 13.1 ± 1.0% (n=7). Average current densities for ASIC3(506-533)1a in the absence (black) and presence (red) of STOM were 69.6 ± 18.7 pA/pF (n=19) and 193.1 ± 61.8 pA/pF (n=18), respectively. FRET efficiency = 1.5 ± 0.5% (n=7). Data given as Mean ± SE. Blue Data point in FRET plots corresponds to control WT ASIC3-CER/STOM-YFP FRET signal replotted from Fig. 3B for comparison.

We then systematically replaced smaller segments of the ASIC3 C-terminus with the corresponding residues from ASIC1a to further narrow down the residues important for the interaction. The precise sequences that are exchanged can be seen in figure 6A. Several of these chimeras exhibited poor expression creating difficulties in selecting cells that consistently exhibited pH-evoked currents. In these cases, we elected to use our fluorescently labelled ASIC3 and STOM for both the functional measurements and the FRET measurements. First, we split the final 28 residues into two chimeras ASIC3(506-519)1a and ASIC3(520-533)1a (Fig. 6B and C). The more proximal chimera, ASIC3(506-519)1a-CER, was still regulated by STOM and was also still able to bind STOM (Fig. 6B). However, the more distal chimera, ASIC3(520-533)1a-CER failed to show any appreciable FRET with STOM-YFP and was not functionally regulated (Fig. 6C). Finally, we made a chimera where the final 8 amino acids of the channel were mutated to their counterparts in ASIC1a, ASIC3(526-533)1a-CER, and both STOM regulation and binding were lost (Fig. 6D). This approach was able to localize a critical binding site on ASIC3 for STOM to the final 8 residues of the channel. Previous work has shown that ASIC3 has a PDZ binding motif in this region that is critical for binding of a number of other proteins including Lin-7B, CIPP, and PSD-95 (Anzai et al., 2002; Eshcol et al., 2008; Hruska-Hageman et al., 2004). Lin-7B and PSD-95 appear to alter ASIC3 surface expression while CIPP shifts the pH dependence of the channel in the basic direction. Interestingly, STOM appears to bind to this same region but lacks a PDZ domain and dramatically reduces ASIC3 currents via an effect on gating.

**Figure 6.**
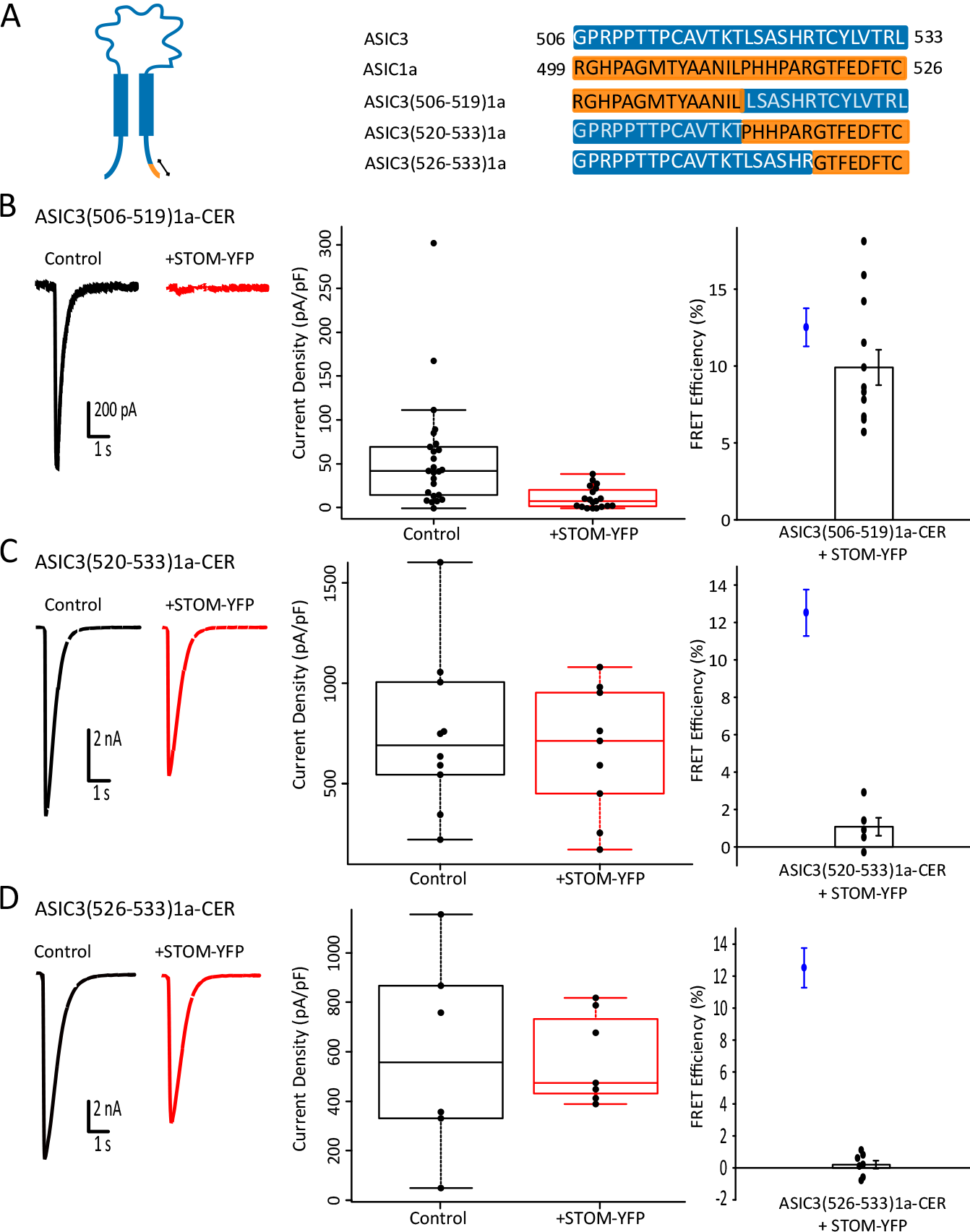
STOM functional interaction of ASIC3 corresponds to binding to the distal C-terminus of ASIC3. (**A**) Cartoon and sequence showing the exact amino acid swap between ASIC3 (blue) and ASIC1a (orange) for the chimeric channels in this figure. (**B-D**) Left shows representative pH 5.5 evoked currents of the indicated chimeric channel alone (black) and co-transfected with STOM-YFP (red). Middle shows boxplots of the current density measurements for the chimeric channel with and without STOM-YFP. Right panel shows the FRET efficiency between STOM-YFP and the indicated ASIC3-CER chimera. Replacement of residues 506-519 of ASIC3 (B) did not alter STOM-YFP binding to, or regulation of ASIC3. However, replacement of residues 520-533 (C) or the even smaller region 526-533 (D) eliminated both regulation and complex formation. Average current densities of ASIC3(506-519)1a-CER in the absence (black) and presence (red) of STOM-YFP were 42.6 ± 6.3 pA/pF (n=23) and 12.0 ± 2.7 pA/pF (n=19), respectively. Average FRET efficiency = 9.9 ± 1.1% (n=12). Average current densities of ASIC3(520-533)1a-CER in the absence (black) and presence (red) of STOM-YFP were 750.1 ± 118.7 pA/pF (n=10) and 661.9 ± 101.2 pA/pF (n=9), respectively. Average FRET efficiency = 1.1 ± 0.5% (n=5). Average current densities of ASIC3(526-533)1a-CER in the absence (black) and presence (red) of STOM-YFP were 586.3 ± 152.7 pA/pF (n=6) STOM-YFP mean 572.9 ± 64.2 pA/pF (n=7), respectively. Average FRET efficiency = 0.2 ± 0.3% (n=7). Data given as Mean ± SE. Blue Data point in FRET plots corresponds to control WT ASIC3-CER/STOM-YFP FRET signal replotted from Fig. 3B for comparison.

### STOM requires TM1 to fully regulate ASIC3

Although swap of distal C-terminus of ASIC3 with the corresponding residues of ASIC1a is sufficient to eliminate STOM functional inhibition, it did not eliminate the possibility that STOM interaction could also involve the N-terminus or TM1. It has previously been reported that N-terminal chimeras between ASIC3 and ASIC1a result in a non-functional channel which we were able to confirm (Salinas et al., 2009). In addition, deletion of portions of the N-terminus yield non-functional channels as well, making this approach a challenge as well (Jasti et al., 2007).

However, simultaneous swap of both the N- and C-termini, has been demonstrated to result in functional ASIC channels (Salinas et al., 2009). Therefore, in order to investigate the role, if any, that the N-terminus of ASIC3 plays in STOM inhibition, we created chimeras that simultaneously swap out the N- and C-terminus of ASIC3. First, creating a chimeric channel with the full N- and C-terminus of ASIC3 replaced with the termini of ASIC1a, ASIC3(1-43,466-533)1a-CER resulted in functional channels (Fig. 7A). As expected, co-expression with STOM-YFP did not affect pH 5.5 evoked currents for ASIC3(1-43,466-533)1a-CER. FRET measurements also showed no signs of interaction between the two proteins. Since the distal C-terminus of ASIC3 was sufficient for STOM-dependent regulation of ASIC3, we inserted this portion of ASIC3 back into the N-C-terminal double chimera. ASIC3(1-43,466-505)1a-CER also yielded functional channels, and STOM-YFP successfully inhibited this chimeric channel (Fig. 7B). Together these data show that the N-terminus of ASIC3 is not a critical region for inhibition by STOM.

**Figure 7.**
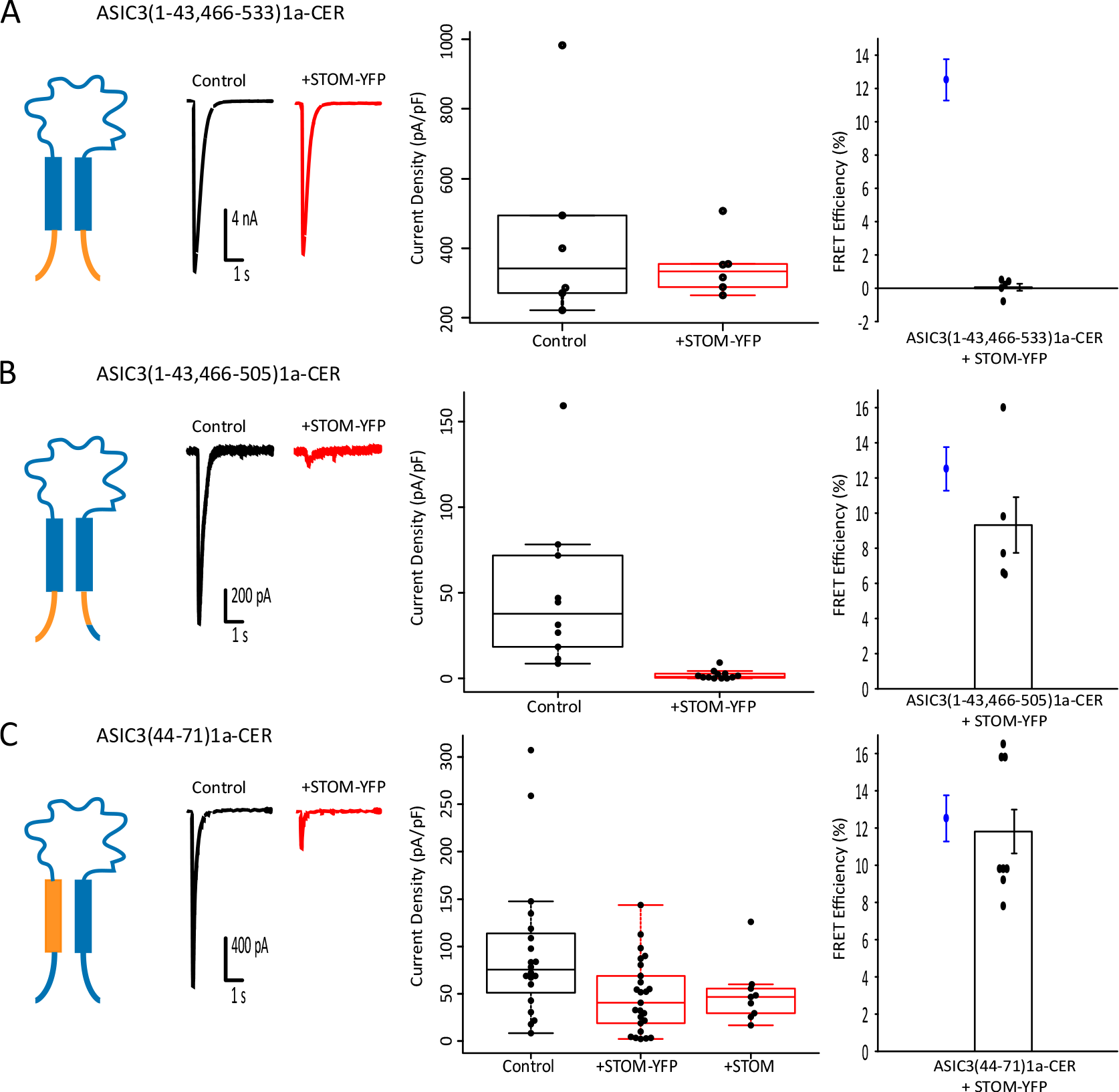
ASIC3’s TM1 but not N-terminus is important for STOM regulation. (**A-C**) Left shows cartoon depicting the chimera being examined and representative pH 5.5 evoked currents of the indicated chimeric channel alone (black) and co-transfected with STOM-YFP (red). Middle shows boxplots of the current density measurements for the chimeric channel with and without STOM-YFP. Right panel shows the FRET efficiency between STOM-YFP and the indicated ASIC3-CER chimera. Replacement of both the N- and C-terminus of ASIC3 with the sequences from ASIC1a (A) eliminates STOM-dependent regulation of ASIC3. However, insertion of the distal C-terminal region back into this N- and C-terminus chimera (B) restored STOM regulation and binding. Swap of the first TM domain of ASIC3 with the first TM of ASIC1a (C) yielded channels whose regulation by STOM was dramatically reduced. Average current densities for ASIC3(1-43,466-533)1a-CER in the absence (black) and presence (red) of STOM-YFP were 334.2 ± 44.2 pA/pF (n=5) and 314.0 ± 15.9 pA/pF (n=5), respectively. Average FRET efficiency = 0.1 ± 0.2% (n=5). Average current densities for ASIC3(1-43,466-505)1a-CER in the absence (black) and presence (red) of STOM-YFP were 37.6 ± 7.9 pA/pF (n=9) and 1.4 ± 0.4 pA/pF (n=11), respectively. Average FRET efficiency = 9.3 ± 1.6% (n=5). Average current densities for ASIC3(44-71)1a-CER in the absence (black) and presence (red) of STOM-YFP or untagged STOM (red) were 72.8 ± 9.0 pA/pF (n=18), 47.3 ± 7.5 pA/pF (n=25) and 40.2 ± 5.0 pA/pF (n=8), respectively. Average FRET efficiency = 11.8 ± 1.2% (n=8). Data given as Mean ± SE. Blue Data point in FRET plots corresponds to control WT ASIC3-CER/STOM-YFP FRET signal replotted from Fig. 3B for comparison.

Lastly, we made a chimeric channel that swapped the TM1 plus a few extracellular residues of ASIC3 for the corresponding regions of ASIC1a. This mutant channel, ASIC3(44-71)1a-CER, was functional, but was only modestly affected by STOM-YFP co-expression (Fig. 7C). Since we showed that the YFP on STOM reduces the overall regulatory effect, we also measured ASIC3(44-71)1a-CER currents in the presence of untagged STOM. In this case ASIC3(44-71)1a-CER currents were decreased only 2-fold as opposed to the nearly 200-fold reduction seen when tagged ASIC3 channels were co-expressed with untagged STOM (Fig. 7C). These data suggest that STOM regulation of ASIC3 is governed by interaction sites on the distal C-terminus and TM1.

## Discussion

Stomatin has been shown to dramatically reduce ASIC3 currents (Brand et al., 2012; Price et al., 2004). In addition, the closely related Stomatin-like (STOML) proteins have been shown to regulate ASICs in an isoform-dependent manner with STOML3 also inhibiting ASIC3 currents (Kozlenkov et al., 2014; Lapatsina et al., 2012; Moshourab et al., 2013; Wetzel et al., 2007). The mechanisms for this inhibition and the binding site for this family of proteins on ASIC3 are not well understood. To answer these questions, we used chimeric channels and combined functional and FRET measurements to localize the binding site for STOM on ASIC3. We made chimeras between the STOM-insensitive ASIC1a and ASIC3. Using this approach, we found that a site that involves the final 8 amino acids of the channel was critical for both binding to and regulation of ASIC3 by STOM, while a second site, on TM1 of ASIC3 was not sufficient for binding, but was necessary for proper regulation of the channel by STOM. In addition, we used confocal microscopy and a surface biotinylation assay to confirm previous findings that STOM did not reduce ASIC3 currents by changing the cell-surface localization of the channel.

Our assay showed no regulation or interaction between ASIC1a and STOM which allowed us to use this chimeric approach. This contrasted with previous work which suggested that ASIC1a bound STOM but was not regulated (Price et al., 2004). The simplest interpretation of our results is that ASIC1a cannot bind STOM which explains the lack of functional regulation. However, FRET requires that the fluorophores be within ~70 Å of each other. It is possible that STOM bound to ASIC1a, but the conformation of the complex is such that the fluorophores are outside the usable range for FRET and that this alternate bound conformation does not result in a regulated channel complex.

There has been very little work examining the binding site between ASIC3 and STOM. However, in one study, the authors posited that a hydrophobic di-peptide (L488 and L489) at the proximal portion of the C-terminus was critical for STOM regulation of the channel (Brand et al., 2012). When these two leucine residues were mutated to aspartate, the effect of STOM was greatly reduced but not eliminated while the binding was unaffected. Our data agree that this region is not critical for the interaction between the two proteins, but do not support this region as critical for STOM-dependent regulation. One possible explanation for the discrepancy in these conclusions stems from the observation that effect of STOM on ASIC3 depends greatly on the expression levels of each protein. In the previous study the LL/DD mutant increased the ASIC3 currents by ~5-10-fold suggesting the possibility that the STOM expression level was not enough to fully inhibit this mutant channel.

Interestingly, the binding site on the distal C-terminus of ASIC3 includes a well characterized PDZ binding motif. This site has been shown to interact with several other PDZ domain containing regulatory proteins including PSD-95, CIPP, and Lin-7b (Anzai et al., 2002; Eshcol et al., 2008; Hruska-Hageman et al., 2004). However, STOM does not contain a PDZ domain and how this interaction occurs is a mystery. Little is known about the targets for SPFH domains, but it is possible that this domain on STOM could be recognizing residues in the distal C-terminus. Further work will be needed to find which domains on STOM are critical for binding to ASIC3. In addition, our data suggests a trend towards decreased ASIC3 currents with just the C-terminal binding site intact (Fig. 7C) which suggests that binding of STOM to the C-terminus might have a modest impact on ASIC3 gating alone.

However, the bulk of the regulatory effect of STOM appears to be mediated through the first transmembrane domain (TM1) of the channel. STOM is an monotopic integral membrane protein that associates with the inner leaflet of the membrane, in part, through a 29 amino acid membrane hairpin. Our data suggest the possibility that the hairpin of STOM could be interacting with and altering the lower portion of TM1. There are a number of residues that are different in this region between ASIC1a and ASIC3. How interaction with this region of the channel could dramatically inhibit gating is unknown. It is possible that the interaction alters the selectivity filter which is thought to be on the intracellular side of TM2 or alters the ability of the TM domains to move during gating (Baconguis et al., 2014; Lynagh et al., 2017; Yoder et al., 2018).

It is also possible that STOM binds specifically to ASIC3 via the C-terminus but alters its function by changing the plasma membrane environment around the channel. STOM is known to bind cholesterol and localize to lipid rafts making it possible that this change in membrane character could explain the change in channel function. Moreover, STOM and the related STOML3 have been hypothesized to impact membrane curvature and stiffness as well which could impact channel function (Brand et al., 2012; Qi et al., 2015). For this hypothesis to be true, the TM1 of ASIC3 would need to be sensitive to changes in the lipid environment in a way that the TM1 of ASIC1a is not.

The physiological role of the ASIC3/STOM complex has not been uncovered to date. However, the closely related STOML3 (60.5% identical, 74.9% similar) has been shown to impact nociceptor mechanosensitivity and be critical for touch sensation (Moshourab et al., 2013; Wetzel et al., 2007). In addition, these studies showed that pH sensitive currents were larger in DRG neurons of STOML3 knockout mice and that the loss of the STOM/ASIC complex can have profound effects on mechanosensitivity in some neurons innervating the skin. We focused this study on the ASIC3/STOM complex, in part, due to the large effect of STOM on ASIC3, but we believe that the results here likely shed light on the regulation of ASIC3 by STOML3 as well.

It is worth wondering about how a complex that renders the channels non-functional might be important for neuronal physiology. We imagine a possible scenario where dynamic regulation of this complex could act to rapidly increase or decrease the amount of functional ASIC3 present on the membrane. There is no direct evidence to date for how this complex might be regulated, but there are palmitoylation sites on STOM that are required for proper membrane targeting of the protein that could be a site of dynamic regulation (Rungaldier et al., 2017). MEC-2, the *C. elegans* homolog of STOM, requires the palmitoylation of a conserved cysteine in order to regulate the ASIC homolog MEC-4/10 (Brown et al., 2008). In addition, the final 30 amino acids of ASIC3 contain six threonines and three serines, including one threonine in the STOM binding site, raising the possibility of a number of potential phosphorylation sites in this region that could lead to dynamic regulation of this complex. A great deal more work needs to be done to uncover the role this complex plays in real cells.

The data presented here show that STOM can inhibit ASIC3 currents by nearly 200-fold. We found two critical sites on ASIC3 for this regulatory effect. The first is a site on the distal C-terminus of the channel that is necessary for both binding between the two proteins and the regulatory effect of the complex. The second critical site on ASIC3 is the first transmembrane domain which appears to be necessary for regulation of the channel by STOM but not for binding. Taken together, we propose a model whereby STOM binds to the distal C-terminus of ASIC3 which leads to a modest reduction in current and anchors the complex together. Then ASIC3 currents are dramatically reduced via a second interaction between TM1 of ASIC3 and the membrane imbedded hairpin of STOM. In order to fully understand this complex, it will be important to understand which parts of STOM are critical for this interaction as well. It will also be interesting to see if the mechanisms that we show here for STOM binding to and regulation of ASIC3 can shed light on the broader question of how this family of proteins regulates such a wide variety of ion channels and transporters.

## Methods

### Mutagenesis

Chimeric channels were created using Gibson Assembly. Sanger sequencing was used to verify correct sequences for all constructs in this study (AGCT, INC., Wheeling, IL USA). For our fluorescently labelled ASIC variants, a short proline rich linker was used to join our fluorophore to the C-terminus of the indicated ASIC isoform. We tried multiple naturally occurring linkers found in the SynLinker database (synlinker.syncti.org) and found that a short linker from the alpha subunit of DNA polymerase worked well (ILPLPYPNSPV) (Liu et al., 2015). Importantly, we found that STOM was unable to regulate ASIC3 when a fluorescent protein was attached directly to the C-terminus of ASIC3. Fluorescently labelled STOM was constructed by joining the fluorophore directly to STOM’s C-terminus. Fluorophores used are indicated throughout and include: mCerulean3 (CER), EGFP (GFP), EYFP (YFP), mTurquoise (TUR), and TagRFP (RFP).

### Cell lines and Transfection

Chinese hamster ovary (CHO-K1) cells (ATCC Manassas, VA USA) were culture in Ham’s F12 media with 10% fetal bovine serum (FBS) at 37°C in 5% CO_2_. Cells at ~70% confluency were transfected via electroporation with a Lonza 4D Nucleofector unit (Lonza, Basel, Switzerland) following the manufacture’s protocol. Plasmid DNA (1 μg) encoding for our WT and mutant rat ASIC3 or ASIC1a proteins in the presence or absence of plasmid DNA (3 μg) encoding for mouse Stomatin was used for transfection. Non-fluorescent ASICs were also transfected with free Citrine plasmid DNA (0.1 μg) to identify cells containing transfected DNA. Following transfection, cells were plated on 12 mm glass coverslips coated in Poly-L-lysine.

### Biotinylation Assay and Western Blotting

Biotinylation of plasma membrane proteins was performed using a slightly modified protocol from a commercially available cell surface protein isolation kit (BioVision, Milpitas, CA USA). CHO-K1 cells were first transfected with ASIC3-TUR or ASIC1a-CER with and without unlabeled STOM. ~18 hrs after transfection, cells were quick-washed in ice-cold PBS followed by incubation with Sulfo-NHS-SS-Biotin at 4°C with gentle shaking. After 1 hr of biotin labelling, reaction was quenched, and cells were scraped and collected followed by centrifugation at 1000X g for 5 min. Cells were washed twice in TBS buffer followed by centrifugation at 1000X g for 5 min. Pellet was collected and cells were resuspended in RIPA buffer for 1 hr at 4°C with end-over-end mixing (in mM): 150 NaCl, 50 TRIS, 1% NP-40, 0.5% sodium deoxycholate, and 1X protease inhibitor added prior to lysis (Thermo Fisher Scientific, Pierce, Rockfrod IL USA). Lysed cells were centrifuged at 10,000X g for 7 min and supernatant was transferred onto packed streptavidin beads and incubated with end-over-end mixing for 1 hr at RT. A portion of each sample was also collected prior to loading onto beads to quantify total protein concentration. Beads were centrifuged at 800X g for 60 sec and supernatant was collected as the non-biotin bound (intracellular) fraction. Beads were washed 3 times in TBS followed by centrifugation at 800X g for 60 sec. Biotin-labelled protein bound to streptavidin beads was eluted by incubating beads with 100 mM DTT for 1 hr at RT. Sample was centrifuged at 800X g for 60 sec and supernatant was collected as biotinylated (plasma membrane) protein.

Samples that were collected just after lysis were measured for the total protein concentration using a BCA assay kit (Thermo Fisher Scientific, Pierce, Rockford, IL USA). Both the biotinylated and intracellular fractions were normalized to total protein and loaded on a 4-12% Bis-Tris pre-cast gel (Thermo Fisher Scientific, Invitrogen, Carlsbad, CA USA) and run at 200 V for 30 min. Protein was transferred from gel to a PVDF membrane at 100 V for 60 min. Membrane was incubated in blocking buffer for 1 hr followed by overnight incubation in primary anti-body (Purified Rabbit anti-GFP, Torrey Pines Biolabs Inc., Secaucus, NJ, USA) at 4°C. Membrane was then washed 6 times with TBS-T followed by 1 hr incubation with secondary antibody (Goat anti-Rabbit IgG, KPL, Gaithersburg, MD USA). Membrane was washed another 6 times in TBS-T then developed in the dark for 5 min with chemiluminescent reagent (Immobilon Forte, Millipore, Burlington, MA USA).

### Electrophysiological Recordings

All experiments were carried out in the whole-cell patch-clamp configuration 24 hr-48 hr post-transfection as described earlier. Borosilicate glass pipettes (Harvard Apparatus, Holliston, MA USA) pulled to a resistance of 2 MΩ - 6 MΩ (P-1000, Sutter Instrument, Novato, CA USA) and filled with an internal solution containing (in mM): 20 EGTA, 10 HEPES, 50 CsCl, 10 NaCl, 60 CsF, with a pH 7.2. Extracellular solution contained (in mM): 110 NaCl, 5 KCl, 40 NMDG, 10 MES, 10 HEPES, 5 Glucose, 10 Trizma Base, 2 CaCl_2_, 1 MgCl_2_, and pH was adjusted as desired with HCl, or NaOH. An Axopatch 200B amplifier and pCLAMP 10.6 (Axon Instruments, Union City, CA USA) were used to record whole-cell currents. Recordings were performed at a holding potential of −80 mV with a 5 kHz low pass filter and sampling at 10 kHz. Channel activation was carried out via a rapid change in solution from a resting pH 8.0 to pH 5.5 for 5 sec. Following activation of the channel by pH 5.5 solution, pH was returned to the resting pH (8.0) for 9 sec and protocol was repeated for a total of six activations. Rapid perfusion was achieved using a SF-77B Fast-Step perfusion system (Warner Instruments, Hamden CT USA). Fluorescence was visualized on an Olympus IX73 microscope (Olympus, Tokyo, Japan) with a CoolLED pE-4000 illumination system (CoolLED, Andover, United Kingdom).

### Förster Resonance Energy Transfer

Cells expressing fluorescent ASIC3 and STOM were examined 16-40 h after transfection using the confocal laser scanning microscope LSM 710 (Zeiss, Thornwood, NY). The same external solutions used for our patch clamp recordings was used for imaging. An area of 500-2500μm^2^ was selected from the overall field of view. Images were taken through a 40× oil objective with a N.A. of 1.3. Cerulean and YFP were excited with separate sweeps of the 458 and 514 nm laser lines of an argon laser directed at the cell with a 458/514 nm dual dichroic mirror. Relative to full power, the excitation power for the imaging sweeps was attenuated to 1% for Cerulean and 0.5% for YFP. Bleaching was performed by using multiple (20-60) sweeps of the YFP laser at full power. Bleaching was usually complete within 30-90s. Emitted light was collected between 460-496nm for Cerulean and 526-579nm for YFP. With this setup, there was no contamination of the relevant Cerulean signal from the YFP. For each experiment, the PMT gain was adjusted to ensure that the maximum pixel intensity was not more than 70% saturated. Fluorescence intensity was then measured by drawing ROIs around the cell in ImageJ (Schneider et al., 2012). Masks were used to eliminate bright fluorescent puncta within the cell. This was a rare occurrence in the Cerulean signal. We also made measurements with ROIs that, to the best of our ability, only surrounded the plasma membrane. This approach did not change the results. FRET efficiency E in percent was calculated as:

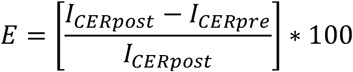

Where I_CERpost_ is the cerulean intensity post-bleaching and I_CERpre_ is the cerulean intensity pre-bleaching.

### Confocal Imaging

ASIC3-TUR and a protein serving as a membrane marker were co-expressed in CHO-K1 cells with and without STOM-YFP and were examined using a Zeiss LSM 710 confocal microscope. The membrane marker protein used was a TagRFP labelled portion of the L-type calcium channel, consisting of the I-II loop of Ca_v_1.2, joined to the N-terminus of Ca_v_1.1 that has previously been shown to associate with the plasma membrane (Kaur et al., 2015; Polster et al., 2018). Excitation and emission for the fluorescent proteins were TUR (ex: 458nm, em: 460-496nm), YFP (ex: 514nm, em: 530-565nm), and tagRFP (ex: 543nm, em: 582-754nm). Relative to full power output, the excitation was attenuated to ~2.5-5% (458 nm), ~5% (514 nm), and ~5-8% (543 nm). Images were obtained with a 40× (1.3 numerical aperture) oil-immersion objective as a single, midlevel optical slice that was halfway between the lower and upper cell surface for CHO-K1 cells.

### Data Analysis and Statistics

Whole cell patch clamp current recordings were analyzed with Clampfit 10.6 software (Axon Instruments, Union City, CA USA). Currents were normalized to cell capacitance and the raw current densities as well as the box plot was plotted for each condition using R software (R Core Team, 2017). All data points collected were plotted, those points that are greater than 1.5 times the boxplot IQR were plotted as outliers. Data reported throughout are calculated as the mean ± the standard error excluding the outliers. Means were also calculated with outliers in table 1 and statistical significance for both conditions were calculated using the Mann-Whitney-Wilcoxon test.

## Acknowledgements

The authors would like to thank Prafulla Aryal for helpful discussions. In addition, we would like to thank Kurt Beam for providing the TagRFP-Ca_V_1.2 I-II loop-Ca_V_1.1 N-Term construct as well as for use of his Zeiss LSM710 confocal microscope. Research reported in this publication was supported by the National Eye Institute of the National Institutes of Health under award number R00EY024267 (to JRB), by the National Institute of Dental and Craniofacial Research under award number F31DE028739 (to MMC), and by the National Heart, Lung, and Blood Institute under award 2T32HL007822 (to RCK). The authors declare no competing financial interest.

---

**Figure 3 - Supplement 1.**
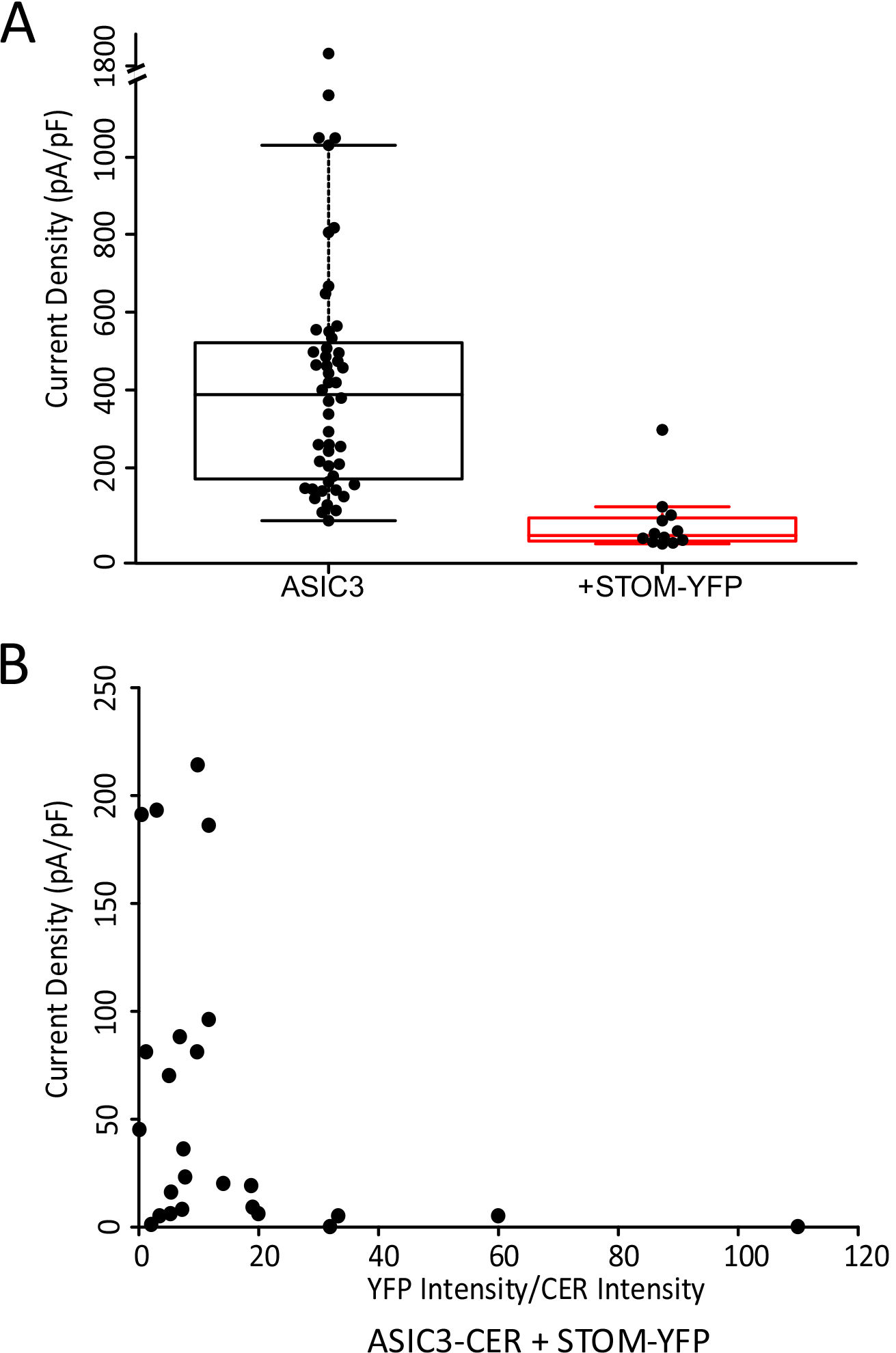
Data examining reduced inhibition with fluorophores. (**A**) Plot of the current densities of the control untagged ASIC3 in the absence (black) (replotted from Fig. 1B) and presence of STOM-YFP Average current densities were 364.0 ± 32.6 pA/pF (n=45) and 34.3 ± 9.1 pA/pF (n=11), respectively. (**B**) Current densities of ASIC3-CER with STOM-YFP from figure 3C plotted as a function of the ratio of fluorescence intensity of YFP:CER. These data show that after about a 15:1 ratio of YFP:CER, the magnitude of the regulation of ASIC3-CER was as large as in the untagged STOM case.

**Figure 4 - Supplement 1.**
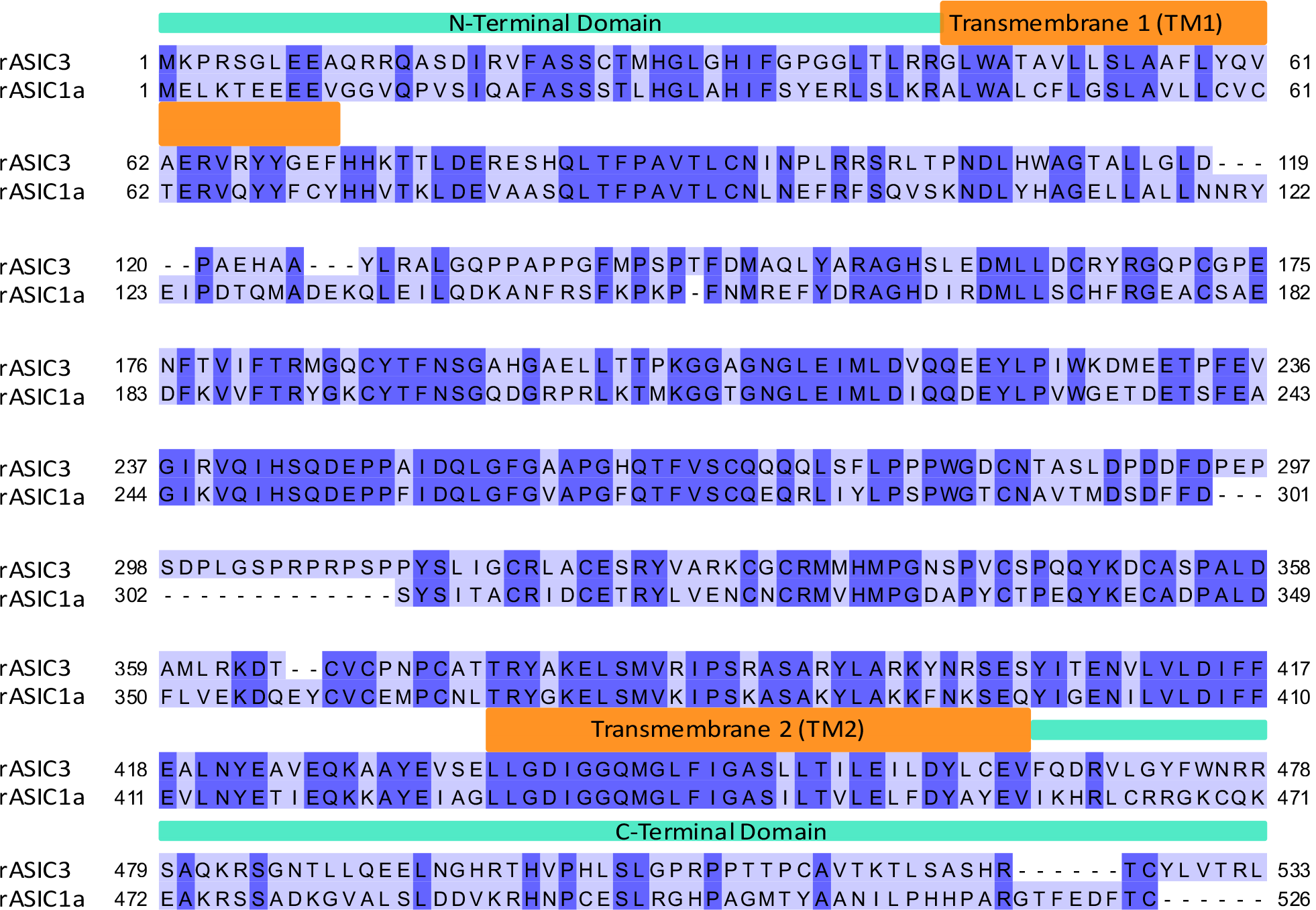
Alignment of rat ASIC3 and rat ASIC1a used for designing chimeras in this study. Dark purple regions show residues that are identical between the two proteins. Orange boxes are present above the residues that make up the transmembrane domains.

**Supplemental Table 1.** Summary of Data

|  | ASIC3 |  | ASIC1a |  | ASIC3-TUR |  | ASIC3-CER |  | ASIC1a-CER |  |
| --- | --- | --- | --- | --- | --- | --- | --- | --- | --- | --- |
| Figure | 1 |  | 1 |  | 2 |  | 3 |  | 3 |  |
| Condition | Control | STOM | Control | STOM | Control | STOM | Control | STOM-YFP | Control | STOM-YFP |
| <u>Including Outliers</u> |  |  |  |  |  |  |  |  |  |  |
| <u>Current Density (pA/pF)</u> | 426.1 | 85.8 | 171.8 | 339.9 | 859.2 | 88.5 | 152.5 | 53.8 | 821.1 | 1227.5 |
| SE | 48.0 | 31.0 | 50.3 | 94.8 | 94.1 | 78.5 | 27.2 | 13.7 | 211.1 | 329.8 |
| n | 48 | 35 | 9 | 9 | 11 | 8 | 23 | 25 | 9 | 9 |
| P |  | 2.05E-09 |  | 0.09 |  | 1.85E-04 |  | 3.7E-04 |  | 0.73 |
| <u>Excluding Outliers</u> |  |  |  |  |  |  |  |  |  |  |
| <u>Current Density (pA/pF)</u> | 364.0 | 2.0 | 122.5 | 200.3 | NA | 4.6 | 137.0 | 47.1 | 509.7 | NA |
| SE | 32.6 | 0.4 | 22.9 | 33.7 |  | 2.7 | 23.5 | 12.4 | 74.2 |  |
| n | 45 | 27 | 8 | 7 |  | 7 | 22 | 24 | 7 |  |
| P |  | 4.9E-13 |  | 0.299 |  | 6.28E-05 |  | 2.6E-04 |  | 0.75 |
| <u>FRET Efficiency (%)</u> |  | NA |  | NA |  | NA |  | 12.5 |  | 1.2 |
| SE |  |  |  |  |  |  |  | 1.2 |  | 0.6 |
| n |  |  |  |  |  |  |  | 11 |  | 8 |
| P (Compared to ASIC3) |  |  |  |  |  |  |  | NA |  | 2.6E-05 |
| P (Compared to ASIC1a) |  |  |  |  |  |  |  | NA |  | NA |

|  | ASIC3(436-533)1a |  | ASIC3(436-465)1a |  | ASIC3(466-533)1a |  | ASIC3(466-505)1a |  | ASIC3(506-533)1a |  |
| --- | --- | --- | --- | --- | --- | --- | --- | --- | --- | --- |
| Figure | 4 |  | 4 |  | 5 |  | 5 |  | 5 |  |
| Condition | Control | STOM | Control | STOM | Control | STOM | Control | STOM | Control | STOM |
| <u>Including Outliers</u> |  |  |  |  |  |  |  |  |  |  |
| <u>Current Density (pA/pF)</u> | 205.3 | 586.8 | 15.4 | 1.6 | 99.5 | 121.4 | 347.2 | 84.3 | 136.4 | 252.4 |
| SE | 50.6 | 257.9 | 4.9 | 1.5 | 26.2 | 41.7 | 124.9 | 67.9 | 41.1 | 82.3 |
| n | 12 | 11 | 9 | 10 | 22 | 9 | 10 | 10 | 22 | 19 |
| P |  | 0.37 |  | 0.001 |  | 0.46 |  | 0.02 |  | 0.47 |
| <u>Excluding Outliers</u> |  |  |  |  |  |  |  |  |  |  |
| <u>Current Density (pA/pF)</u> | NA | 337.1 | NA | 0.0 | NA | NA | 239.9 | 13.4 | 69.6 | 193.1 |
| SE |  | 109.0 |  | 0.0 |  |  | 80.3 | 5.1 | 18.7 | 61.8 |
| n |  | 10 |  | 9 |  |  | 9 | 9 | 19 | 18 |
| P |  | 0.57 |  | 4.1E-04 |  | NA |  | 0.02 |  | 0.29 |
| <u>FRET Efficiency (%)</u> |  | NA |  | NA |  | 1.0 |  | 13.1 |  | 1.5 |
| SE |  |  |  |  |  | 0.5 |  | 1.0 |  | 0.5 |
| n |  |  |  |  |  | 8 |  | 7 |  | 7 |
| P (Compared to ASIC3) |  |  |  |  |  | 2.6E-05 |  | 0.58 |  | 6.3E-05 |
| P (Compared to ASIC1a) |  |  |  |  |  | 0.85 |  | 3.1E-04 |  | 0.48 |

**Supplemental Table 1 Continued.**
|  | ASIC3(506-519)1a-CER |  | ASIC3(520-533)1a-CER |  | ASIC3(526-533)1a-CER |  | ASIC3(1-43,466-533)1a-CER |  | ASIC3(1-43,466-505)1a-CER |  |
| --- | --- | --- | --- | --- | --- | --- | --- | --- | --- | --- |
| **Figure** | 6 |  | 6 |  | 6 |  | 7 |  | 7 |  |
| **Condition** | Control | STOM-YFP | Control | STOM-YFP | Control | STOM-YFP | Control | STOM-YFP | Control | STOM-YFP |
| <u>Including Outliers</u> |  |  |  |  |  |  |  |  |  |  |
| <u>Current Density (pA/pF)</u> | 58.1 | 12.0 | 750.1 | 661.9 | 586.3 | 572.9 | 442.3 | 346.0 | 49.8 | 2.0 |
| SE | 12.6 | 2.7 | 118.7 | 101.2 | 152.7 | 64.2 | 105.3 | 32.1 | 13.5 | 0.7 |
| n | 25 | 19 | 10 | 9 | 6 | 7 | 6 | 6 | 10 | 12 |
| P |  | 1.4E-04 |  | 0.3 |  | 0.84 |  | 1 |  | 1.1E-04 |
| <u>Excluding Outliers</u> |  |  |  |  |  |  |  |  |  |  |
| <u>Current Density (pA/pF)</u> | 42.6 | NA | NA | NA | NA | NA | 334.2 | 314.0 | 37.6 | 1.4 |
| SE | 6.3 |  |  |  |  |  | 44.2 | 15.9 | 7.9 | 0.4 |
| n | 23 |  |  |  |  |  | 5 | 5 | 9 | 11 |
| P |  | 3.3E-04 |  | NA |  | NA |  | 1 |  | 1.9E-04 |
| <u>FRET Efficiency (%)</u> |  | 9.9 |  | 1.1 |  | 0.2 |  | 0.1 |  | 9.3 |
| SE |  | 1.1 |  | 0.5 |  | 0.2 |  | 0.2 |  | 1.6 |
| n |  | 12 |  | 5 |  | 7 |  | 5 |  | 5 |
| P (Compared to ASIC3) |  | 0.1 |  | 4.6E-04 |  | 6.3E-05 |  | 4.6E-04 |  | 0.11 |
| P (Compared to ASIC1a) |  | 2.5E-04 |  | 0.97 |  | 0.24 |  | 0.14 |  | 0.002 |

|  | ASIC3(44-71)1a-CER |  |  | ASIC3 |  |
| --- | --- | --- | --- | --- | --- |
| **Figure** | 7 |  |  | 3-Supplement 1 |  |
| **Condition** | Control | STOM-YFP | STOM | Control | STOM-YFP |
| <u>Including Outliers</u> |  |  |  |  |  |
| <u>Current Density (pA/pF)</u> | 93.8 | 47.3 | 49.7 | 426.1 | 56.3 |
| SE | 16.4 | 7.5 | 10.0 | 48.0 | 22.6 |
| n | 20 | 25 | 9 | 48 | 12 |
| P |  | 0.013 | 0.045 |  | 1.2E-06 |
| <u>Excluding Outliers</u> |  |  |  |  |  |
| <u>Current Density (pA/pF)</u> | 72.8 | NA | 40.2 | 364.0 | 34.3 |
| SE | 9.0 |  | 5.0 | 32.6 | 9.1 |
| n | 18 |  | 8 | 45 | 11 |
| P |  | 0.04 | 0.024 |  | 5.4E-07 |
| <u>FRET Efficiency (%)</u> |  |  | 11.8 |  | NA |
| SE |  |  | 1.2 |  |  |
| n |  |  | 8 |  |  |
| P (Compared to ASIC3) |  |  | 0.7 |  |  |
| P (Compared to ASIC1a) |  |  | 1.6E-04 |  |  |

